# The mechanistic basis of evolutionary transitions between grey, slate, and blue colour in Tanagers (Thraupidae)

**DOI:** 10.1101/2023.10.31.564904

**Authors:** Frane Babarovic, Christopher R Cooney, Thomas Zinn, Szymon M. Drobniak, Katarzyna Janas, Jonathan D Kennedy, Nicola J Nadeau, Andrew J Parnell, Gavin Huw Thomas, Stephanie Burg

## Abstract

Both pigmentary and structural colours share many common elements of their feather anatomy, i.e. keratin, air and melanin packed in the melanosomes, despite utilizing different mechanisms of the colour production. This means that evolutionary transitions between pigmentary and structural colours can be achieved through a simple adjustment of these elements. Recently, an evolutionary hypothesis for the transition between pigmentary grey, through slate and finally to structural blue colour has been proposed and confirmed in the clade Tanagers on a macroevolutionary level. Here, we investigate mechanistic basis of this evolutionary pathway. By using SAXS (small-angle X-ray scattering) we have quantified important elements of spongy layer in medullary cells that is crucial for colour production by coherent scattering of light wavelengths. We have quantified five elements of the spongy layer: nanostructure complexity, average hard block thickness, average soft block thickness, filling fraction and I_o_ value. We report that across different categories of feather colour, i.e. blue, slate and grey, nanostructure complexity, filling fraction and I_o_ value explained variation in the chromatic component of the colour (between the three colour categories). Chromatic variation within the colour category was explained by filling fraction in the case of slate colour and by nanostructure complexity and average hard block thickness in the case of blue colour. We propose that variation in different elements or combination of elements of the spongy nanostructure has been utilised in feather colour evolution, both within and between colour categories, to overcome developmental constraints imposed by self-assembly processes.

## Introduction

Birds are one of the most colourful groups of animals (Cuthill et al., 2017). The mechanisms by which they achieve their full colour gamut range from structural to pigmentary as well as the combination of both (Shawkey & D’Alba, 2017; Stoddard & Prum, 2011). The breadth of the plumage colour spectrum relies on the internal architecture of feathers (either variation in feather nanostructure and/or pigment composition) in both types of colour producing mechanisms (Prum, 2006; McGraw, 2006). Therefore, to understand the evolution of plumage colouration, it is critical to study the elements of feather nanostructure that participate in colour production (Maia et al., 2013).

Pigmentary colours are produced by selective absorption and reflection of certain wavelengths of light from the pigments deposited in feathers and the colour produced will depend on the chemical composition of the pigments themselves (McGraw, 2006). The most common pigments in birds’ plumage are melanins (brown, grey and black colour) and carotenoids. In melanin-based plumage colouration, melanin is stored within melanosomes, which are organelles that produce, transport and store melanin pigment (Marks & Seabra, 2001; D’Alba & Shawkey, 2019). It has been shown that different melanosome shapes are characteristic of different melanin-based plumage colouration (Babarović et al., 2019; Li et al., 2010; Nordén et al., 2019). For example, grey plumage colouration has characteristic melanosomes that are larger than any other melanosomes in pigmentary melanin colouration (Babarović et al., 2019; Li et al., 2010). The concentration of melanosomes is also important for melanin-based pigmentary colours with increasing concentration contributing to darker colours (Field et al., 2013).

In structural colour, the colour is produced by coherent scattering of light as it interacts with the interface of nanoscale structures within the feathers, normally biopolymer (chitin and beta-keratin) and air that possess different refractive indices (Burg & Parnell, 2018; Prum, 2006). In iridescent structural colours in feathers, the colour producing nanostructure consists of a periodical arrangement of melanosomes embedded in keratin on the periphery of the feather barbules (Prum, 2006). Colours produced in this way are angle dependent (changing hue with the changing viewing angle) (Kinoshita et al., 2008; Nordén et al., 2021). In contrast, non-iridescent structural colours in feathers, are independent of viewing angle, and are often purple, blue and UV in hue (Prum, 2006; Fan et al., 2019). In these instances, the colours are produced by coherent scattering of light by the nanoscale arrangement of keratin and air in the medullary cells of feather barbs. A keratin matrix is placed above this nanostructure (towards the edge of the feather barbs) while a layer of melanosomes is located below it (i.e. towards the central shaft of feather barbs) (Fan et al., 2019; Prum, 2006; Shawkey et al., 2003; Shawkey & Hill, 2006). In addition, characteristics of the melanosomes (size and shape) are also correlated with structural colours (Babarović et al., 2019; Li et al., 2010). For example, melanosomes found in non-iridescent structural colours are bigger than in most other colour categories and they overlap in shape with melanosomes characteristic for grey pigmentary colour (Babarović et al., 2019).

In non-iridescent structural colour production, keratin and air are structured in the medullary cells and this can be ordered in two possible ways to produce coherent scattering and ultimately colour production (Prum, 2006; Saranathan et al., 2012). Sphere type nanostructure consists of numerous spherical air cavities uniform in their length scale and interconnected by small air passages that are embedded in the keratin matrix. Channel type nanostructure consists of elongated and often rotated air channels embedded in a keratin matrix that creates keratin bars around them. In both nanostructure architectures, there is a periodicity between the two different refractive indices, with a length scale on the order of the wavelength of visible light which produces coherent scattering (Prum, 2006; Prum et al., 2009; Saranathan et al., 2012). In this type of scattering, colour isproduced as a sum of the interactions among scattered waves (Prum et al., 1998). Variation in the physical parameters of the nanostructure, as well of the other components of the barb (the thickness of the keratin matrix as well as melanosomes layer), will influence the hue of the producedcolour. Namely, uniformity of the diameter of keratin rods strongly predicts spectral saturation while chromatic variation is related to the spatial frequency and thickness of the spongy layer, the ratio of the amount of spongy layer to melanin and the thickness of keratin layer above the spongy layer (Fan et al., 2019; Shawkey et al. 2003). Therefore, colour variation in non-iridescent structural colours is not produced by absence or presence of any of these structural elements, but rather bythe difference in their properties.

Despite the differences in colour production mechanisms, feathers exhibiting melanin-based pigmentary colours and structural colours in many cases have similar building materials, i.e. keratin and melanin packed in melanosomes (McGraw, 2006; Prum, 2006; Shawkey & D’Alba, 2017). This similarity in structural components has led to the hypothesis that evolutionary transitions between pigmentary and structural colours in birds’ plumage can proceed through structural rearrangement of already pre-existing elements within the feathers, rather than evolution of a completely novel phenotype (Prum, 2006, Shawkey et al., 2006). This is referred to as ‘evolutionary tinkering’ toreflect the idea that modifications of an existing phenotype can lead to a novel phenotype (Bockaert & Pin, 1999; Jacob, 1977; Saraste & Castresana, 1994). This type of evolutionary transition has already been detected in birds’ plumage (Shawkey et al., 2006; Driskell et al., 2010; Doucet et al., 2004). For example, evolutionary transitions between matte black plumage and iridescent plumage colouration in grackles and allies depend on rearrangement of melanosomes (Shawkey et al., 2006). In feathers with matte black plumage, melanosomes are scattered evenly around barbules while in iridescent feathers melanosomes are arranged in layers near the edges of the barbules (Shawkey et al., 2006). This ordering of melanosomes creates interfaces with beta keratin and is responsible for coherent scattering and therefore colour production.

Recently, it has been proposed that grey (a pigmentary colour) and blue (a non-iridescent structural colour) are evolutionarily linked (Babarović et al., 2019). For a phylogenetically wide range of feathers, an investigation of the shape of the melanosomes placed underneath the spongy layer revealed that they overlap in shape with the melanosomes characteristic of grey pigmentary feathers (Babarović et al., 2019). Furthermore, rudimentary spongy nanostructure, whose colouration has been described as slate (grey-blue or blue-grey), was proposed to be an intermediary link between pigmentary grey and structural blue colour (Saranathan et al., 2012). Finally, recently, a macroevolutionary transition between these colours has been confirmed in the Tanager clade (Aves: Thraupidae) (Babarović et al., 2023). In Tanagers, transitions between grey and slate were found to be common, but blue colour was found to evolve only from the slate colour. Nevertheless, a mechanical basis of these evolutionary transitions has not been tackled previously. Specifically, we do not know what structural elements of the spongy structure in feather barbs are changing to enable this transition.

Here, we investigated the nanostructural characteristics of elements of the medullary (or spongy) layer in blue, slate and grey feathers, i.e. air and keratin matrices, in Tanagers (Aves: Thraupidae). Our research is focused on the chromatic variation of the colour, i.e. hue and saturation, across blue,slate and grey colour categories. The Tanagers are large radiation of birds with a primarily Neotropical distribution and a diverse array of plumage colours including many species with blue, slate, and grey plumage colour. We used small angle x-ray scattering (SAXS) to assess several nanostructural elements of grey, slate and blue feathers in Tanagers to understand: i) what structural elements are responsible for the colour differences between these three colour categories? and ii) what structural elements account for colour variance within slate and blue colour categories?

## Materials and methods

### Feather sampling

We sampled feathers at the Zoological Museum, Natural History Museum of Denmark, University of Copenhagen. We sampled 10 species for grey feathers, 16 species for slate feathers and 11 species for blue feathers. Across all species, we sampled from following patches: wing covert, breast, nape, rump, throat, and mantle. We aimed at sampling one feather from three different bird skins fromthe same plumage patch. In total, 117 feather samples were collected (30 grey feathers, 48 slate and33 blue feathers). (Full report on sampling details are in Supplement material: Table S1). Feather sampling was designed to ensure coverage of a wide range of the grey, slate and blue colour gamut and was informed by analysis of colour categorization from written descriptions of plumage colouration from Birds of the World and digitally calibrated images of plumage colours in Tanagers (Babarović et al., 2023: Distinctiveness analysis; Billerman et al., 2022).

### Reflectance data

The reflectance of each collected feather was measured using an Ocean Optics USB2000+ spectrometer with UV transmissive fibre optic cable. A Y-shaped cable was connected to the light source, spectrometer and a third opening was mounted to the sample. The light source used was A DT-MINI-2-GS (Ocean Optics) Deuterium Tungsten Halogen UV-Vis-NIR light source with wavelength range from 215-2500 nm. The probe was placed 5 mm from the feather sample at 90 degrees to produce a small spot of light (∼ 1 mm in diameter). To maximise the reflectance signal as much as possible, we populated the ∼1 mm light spot with as many distal and coloured contour feather barbs as possible (∼3 barbs). The measurements were acquired with the Spectra Suite (Ocean Optics) software with an integration time of 300 ns, 3 scans to average and 3 nm boxcar width. Thecollected reflectance spectra were then normalized by dividing the results by the spectra collected from a white standard (a Polytetrafluoroethylene (PTFE) diffuse white standard (Labsphere)) measured under the same instrumental conditions.

Spectral data were further analysed in R using the package “pavo” (version 2.7.1) (Maia et al., 2019; R Core Team, 2021). Spectra were first individually smoothed and then averaged on a species level (measurements from three feathers were averaged) with “Procspec” and “aggspec” functions, respectively. Next, we estimated the chromatic properties (hue and saturation) of the measured spectra by estimating avian cone catch values (u, s, m, l) associated with each spectrum using the “vismodel” function. The UVS avian visual system was used as the visual model since genomic sequencing of the UV/violet SWS1 cone opsin gene indicated the presence of amino acid residues signifying UV sensitivity in Tanagers (Ödeen & Håstad, 2013).

### Small Angle X-ray Scattering (SAXS)

SAXS data for the spongy layer in the medullary cells of the feather barbs were collected at the Diamond Light Source (UK) with the beamline I22. Historically, the internal structure of feathers has been investigated using different microscopy techniques, with Transmission Electron Microscopy (TEM) yielding most detailed results. Limitations, however, do exist with the TEM approach. Namely, artificial shrinkage of the samples during the sample preparation as well as time-consuming sample preparation. In contrast, SAXS requires no sample prep, beyond mounting the sample in the path of the beam (Saranathan et al., 2012; Janas et al., 2020; Parnell et al., 2015).

SAXS was performed on the samples mounted over 3mm apertures on an aluminium sample plate perpendicular to the direction of the x-rays. Scattering of the photons occurs at interfaces in the biological material, here the electron density contrast produces a diffraction pattern that is detected by a 2-D detector. In the case of colour producing nanostructures in feather barbs, the diffraction pattern will take a circular form due to the isotropic nature of the structure. The data is reduced to a 1D scattering pattern by radially integrating the 2D detector image with I (intensity) on the y-axis and q (scattering vector) on the x-axis. Bright rings in the diffraction pattern will correspond to a peak in the 1D scattering profile. In samples which lack colour-producing nanostructure in the feather barb, the scattering plot will be featureless with no peaks detected (Saranathan et al., 2012; Prum et al., 1998). At Diamond, an x-ray wavelength of 1.2 Å (10 keV) was used with a rectangular shaped microfocus beam (20 μm x 20 μm) and a Pilatus P3-2M 2D detector placed at the 9.575 m from the sample. This setup allowed a length scale of 620 nm as an upper resolution.

We aimed to scan the same regions of the feather using SAXS as were measured for the spectrometer measurements. For each barb scanned (117 in total), either 121 or 49 individual 2D SAXS images were collected (frames) using a raster scan. For each measured frame a scattering profile with intensity (I) as a function of q (scattering wavevector q=4νSin8/τχ) was extracted withthe DAWN software (Filik et al., 2017). Following this, for each feather, we calculated the sum value in intensity (I) for each scattering profile and selected the top 3 scattering profiles with the highest summed scattering intensities. This resulted in a total of 351 scattering profiles, i.e. three for each of the 117 feathers which were carried forward for 1) peak and shoulder detection analysis and 2) One-dimensional correlation function analysis (CORFUNC) (Strobl & Schneider, 1980). The analysis was implemented in the custom python code, written by Dr Adam Washington, and modified for the purpose of this research by Dr Stephanie Burge.

## Analysis

### Principal component analysis

We transformed the reflectance spectra measurements into cone catch values (u, s, m and l) which estimate the chromatic properties of colour (hue and saturation), as birds see them (Stoddard & Prum, 2008). Cone catch values describe a point in the colourspace, a morphospace adjusted to ultraviolet-sensitive avian visual system (Ödeen & Håstad, 2013; Stoddard & Prum, 2008). Furthermore, we used Principal Component Analysis (PCA; Jolliffe, 2002) to reduce the dimensionality of the colourspace. Therefore, the principal components capture both elements of the chromatic variation (hue and saturation) of the measured colour.

### Peak and shoulder detection analysis

Every SAXS profile of a feather containing nanostructure will contain 1) shoulders, 2) peaks or 3) both (explained further down) (Saranathan et al., 2012). If the nanostructure responsible for the structural colour is absent, the scattering intensity will decrease with increasing q (spatial frequency of variation in electron density) with no detectable features (Fig. 1, a). In the scattering patterns, a shoulder without any peaks represents a feather with a rudimentary spongy layer in the medullary cells of the feather barbs, this is a structure organized enough to produce coherent scattering and therefore structural colour, but not sufficiently monodisperse to generate a sharp peak (Fig. 1, b). In contrast, a peak in the scattering pattern represents a feather where the medullary cells in the feather barbs have short-range periodicity in the spongy layer and a more uniform length scale distribution resulting in a more well-defined scattering feature (Fig. 1, c). Furthermore, additional peaks and/or shoulders detected after the first peak demonstrates a long-range periodicity in the nanostructure not present in a nanostructure with just one peak/shoulder (Fig. 1, c-d). The numberof higher order features corresponds to the number of elements following peak or a shoulder (more than one scattering feature) (Fig. 1, c-d). Any scattering pattern with just one peak or one peak andadditional shoulders represents channel-type spongy layer (Fig. 1, c) while patterns with additional peaks after the first peak is representative of sphere-type nanostructure in the spongy layer (Fig. 1,d).

**Figure 1.**
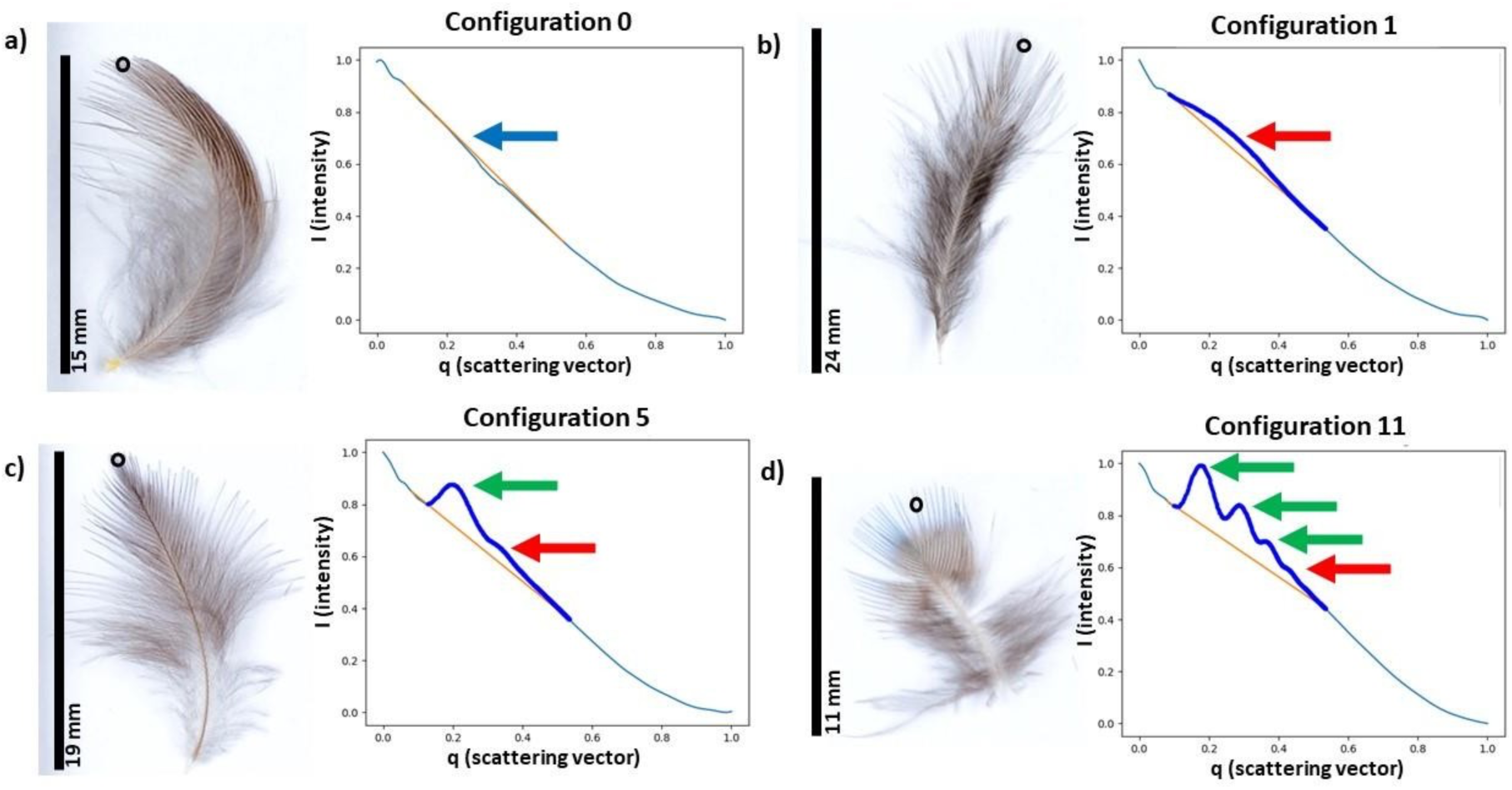
Examples of scattering profiles and feathers where the measurements were taken. On each panel first image represents the feather and second an accompanying scattering profile. For each panel, the SAXS measurement is taken on the spot marked with the black circle on the feather. Features describing nanostructure components in each scattering panel are marked with arrows: blue arrow represents lack of the scattering feature, red arrow represents shoulder, and green arrow represents peak. The figure is a visual representation of Table 2, and the classification of combinations of features is explained in the table. Scattering profiles of other possible configurations are represented in the Supplement material Figure S1. Panel (a) represents configuration 0, with a lack of any structural components. Feather where this scattering plot was obtained is from is the mantle of Double-collared seedeater (*Sporophila caerulescens*). Panel (b) represents configuration 1, with a one shoulder detected and is typical for the rudimentary form of the spongy nanostructure in the medullary feather cells. Feather where this scattering plot was obtained is fromis the rump of Black-throated flowerpiercer (*Diglossa brunneiventris*). Panel (c) representsconfiguration 5, with a one peak and one shoulder detected and is typical for the channel-type spongy layer. Feather where this scattering plot was obtained is from is the rump of Masked flowerpiercer (*Diglossa cyanea*). Panel (d) represents configuration 11, with three peaks and one shoulder detected and is typical for the sphere-type spongy layer. Feather where this scattering plot was obtained is from is the breast of Blue dacnis (*Dacnis cayana)*.

To detect and classify these features in the 351 scattering patterns, we developed code in Python to detect peaks and shoulders. Peaks were defined as a point where the derivative of the 1D curve was equal to 0 and the second derivative was negative (Stewart, 2005). In each instance that a peak was detected, a Gaussian curve was fitted to the local peak which returned the peak intensity (I_o_), the peak position (q_m_), and the standard deviation or “width” (σ) of the peak (Additional table 4.; https://figshare.com/s/1110fce894e65a69c329) (Stewart, 2005). For shoulder detection, we used the “Kneedle” approach which searches for a point of maximum curvature in the function defined as a peak in a calculated detection function based on the sum of the vertical and perpendicular distance between the function and a straight line (Satopaa et al., 2011). When the algorithm detects a shoulder is it is characterized by a (I_o_, q_m_) value indicating this point of maximum curvature (Table S4.). The max I_o_ value of the first feature detected in the scattering plots where nanostructure is present is proportional to the thickness of the spongy layer in the medullary cell. The q_m_ position corresponds to the dominate lengthscale or spacing within the nanostructure calculated in as 2π/q_m_. We used I_o_ for the further analysis by choosing the value of the I_o_ for each species of the highest average values across 3 feathers (Supplement material: Table S3.)

Examining our results, the possible scattering patterns across all the feathers had a limited number of peak and shoulder configurations. A scoring system for the scattering patterns was used to classify and sort these configurations as follows: i) peak is scored as 3, ii) shoulder after the peak is scored as 2, and iii) just a shoulder is scored as 1. The highest scoring nanostructure is 13 with three peaks and two shoulders (Fig. 1, d), while the lowest is zero with no nanostructure detected (Fig. 1, a). We termed this variable “nanostructure complexity” and used it for further analysis. Nanostructure complexity indicates a length-scale of periodicity with higher values indicating nanostructures with a longer-range periodicity than smaller values. Due to our scoring system, some configurations are not possible, i.e. nanostructure scoring of 4, 7, 9 and 12. The scoring system, all possible configurations, and their meanings are reported in the Table 1 and Supplement material: Figure S1. The representative of the main configuration and the feathers from which the measurements were taken are illustrated in the Fig. 1. The scores are reported in Supplement material: Table S4. For species level score of the nanostructure, a highest score of the nanostructure among 9 frames from 3 featherswas taken (Additional table 4.; https://figshare.com/s/1110fce894e65a69c329).

**Table 1.**
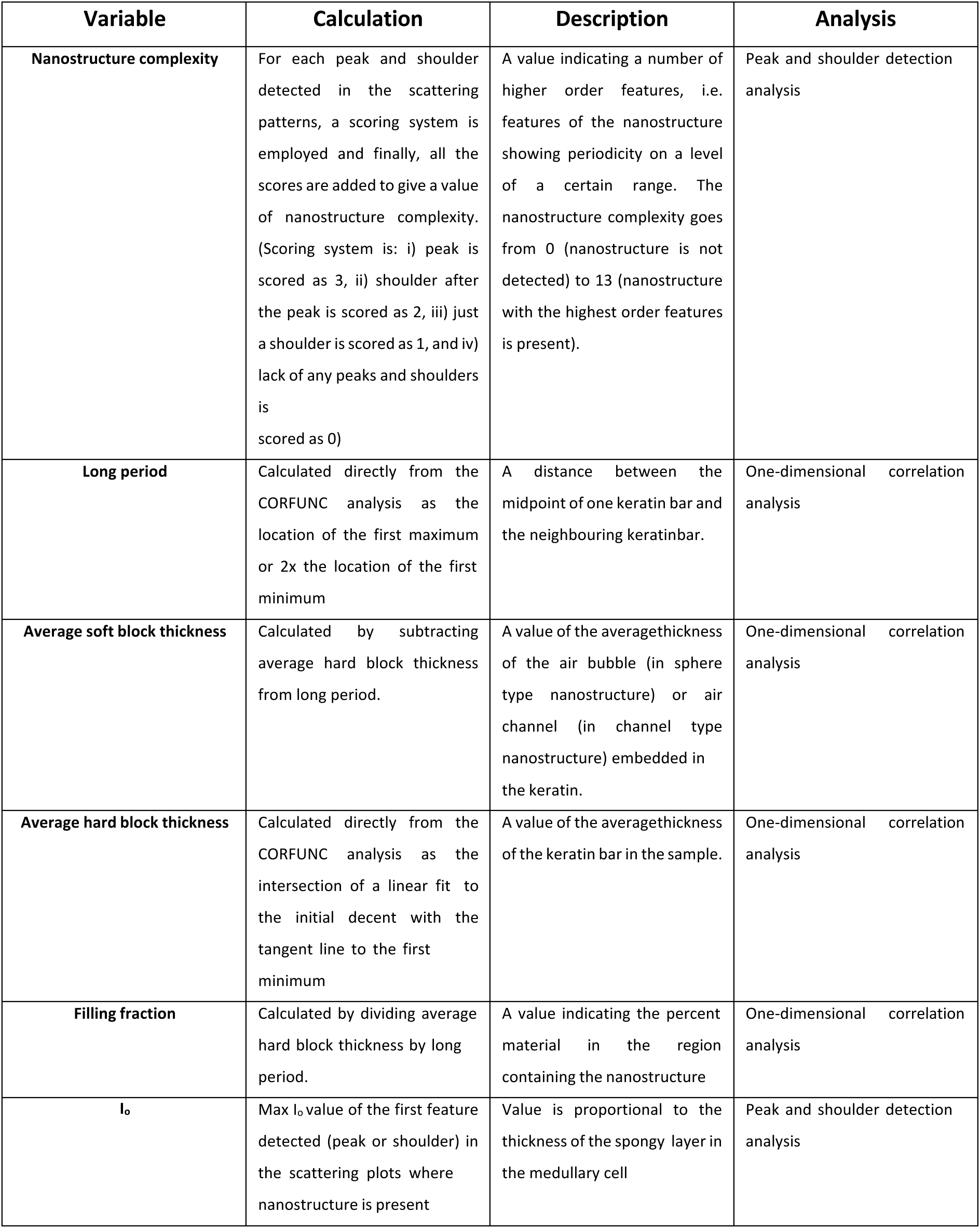
Variables extracted from the Peak and shoulder detection analysis and One-dimensional correlation analysis of the Small-angle X-ray scattering experiment. For each variable (first column), a description of how the variable is calculated (second column), what part of the nanostructure it quantifies (third column) and which analysis is used to obtain the variable (fourth column) is listed.

| Variable | Calculation | Description | Analysis |
| --- | --- | --- | --- |
| **Nanostructure complexity** | For each peak and shoulder detected in the scattering patterns, a scoring system is employed and finally, all the scores are added to give a value of nanostructure complexity. (Scoring system is: i) peak is scored as 3, ii) shoulder after the peak is scored as 2, iii) just a shoulder is scored as 1, and iv) lack of any peaks and shoulders is scored as 0) | A value indicating a number of higher order features, i.e. features of the nanostructure showing periodicity on a level of a certain range. The nanostructure complexity goes from 0 (nanostructure is not detected) to 13 (nanostructure with the highest order features is present). | Peak and shoulder detection analysis |
| **Long period** | Calculated directly from the CORFUNC analysis as the location of the first maximum or 2x the location of the first minimum | A distance between the midpoint of one keratin bar and the neighbouring keratinbar. | One-dimensional correlation analysis |
| **Average soft block thickness** | Calculated by subtracting average hard block thickness from long period. | A value of the averagethickness of the air bubble (in sphere type nanostructure) or air channel (in channel type nanostructure) embedded in the keratin. | One-dimensional correlation analysis |
| **Average hard block thickness** | Calculated directly from the CORFUNC analysis as the intersection of a linear fit to the initial decent with the tangent line to the first minimum | A value of the averagethickness of the keratin bar in the sample. | One-dimensional correlation analysis |
| **Filling fraction** | Calculated by dividing average hard block thickness by long period. | A value indicating the percent material in the region containing the nanostructure | One-dimensional correlation analysis |
| **$I_0$** | Max $I_0$ value of the first feature detected (peak or shoulder) in the scattering plots where nanostructure is present | Value is proportional to the thickness of the spongy layer in the medullary cell | Peak and shoulder detection analysis |

**Table 2.** Overview of the nanostructure complexity variable. The first column lists all the possible values of the variable. Column two shows absence (first row) and presence (the rest of the rows) and the count of structural elements for each score of the nanostructure complexity. Values of the nanostructure complexity are calculated by addition of the scores associated with each structural elements detected for each category. Scoring system is as follows: i) peak is scored as 3, ii) shoulder after the peak is scored as 2, and iii) just a shoulder is scored as 1. Column three shows the biological meaning of every score of nanostructure complexity. In short, score 0 indicates no nanostructure detected, scores 1 – 2 indicate rudimentary nanostructure, scores 3 – 5 show channel-type nanostructure and finally, scores 6 – 13 indicate sphere-type nanostructure. Columns four, five and six show the number of feathers and species where each nanostructure complexity score wasdetected across grey, slate and blue colour category.

| Nanostructure complexity | Elements detected | Biological meaning | Grey colour | Slate colour | Blue colour |
| --- | --- | --- | --- | --- | --- |
| 0 | 0 peaks, 0 shoulders | No Nanostructure | 25 feathers from 9 species | 8 feathers from 4 species | 0 feathers |
| 1 | 0 peaks, 1 shoulder | Rudimentary nanostructure | 2 feathers from 1 species | 26 feathers from 13 species | 0 feathers |
| 2 | 0 peaks, 2 shoulders |  | 0 feathers | 5 feathers from 2 species |  |
| 3 | 1 peak | Channel-type nanostructure | 0 feathers | 3 feathers from 1 species | 5 feathers from 2 species |
| 4 | Not possible |  |  |  |  |
| 5 | 1 peak, 1 shoulder |  | 0 feathers | 2 feathers from 1 species | 12 feathers from 5 species |
| 6 | 2 peaks | Sphere-type nanostructure | 0 feathers | 0 feathers | 3 feathers from 2 species |
| 7 | Not possible |  | 0 feathers | 0 feathers |  |
| 8 | 2 peaks, 1 shoulder |  | 2 feathers from 1 species | 4 feathers from 3 species | 9 feathers from 4 species |
| 9 | Not possible |  |  |  |  |
| 10 | 2 peaks, 2 shoulders |  | 1 feather from 1 species | 5 feathers from 3 species | 1 feather from 1 species |
| 11 | 3 peaks, 1 shoulder |  | 0 feathers | 0 feathers | 6 feathers from 2 species |
| 12 | Not possible |  |  |  |  |
| 13 | 3 peaks, 2 shoulders |  | 0 feathers | 2 feathers from 1 species | 0 feathers |

### One-dimensional correlation analysis

To extract length scale values of the nanostructure elements in the medullary cells spongy layerfrom the SAXS scattering profiles we used a one-dimensional correlation analysis known as CORFUNC (Strobl & Schneider, 1980). The foundation of this analysis is a Fourier transform of the 1-dimensional scattering profiles with the assumption that the system is a two-phase system of different electron densities. In our case this is keratin and air. The analysis involves extrapolating the low-q scattering data to a zero by fitting it to a Guinier curve and extrapolating the high-q scattering data to infinity using a Porod curve (Strobl & Schneider, 1980). The experimental data together with the extrapolated data across the new q range (from zero to infinity) is then Fourier transformed and returns the real space correlation function for the feather specimen. Finally, a linear fit together with the position of the first minimum and first maximum of the correlation function is used to extract the length scales of elements of the medullary cells spongy layer based on a two-phase assumption.

Therefore, for further analysis, we have extracted the following values: 1. Average hard block thickness – a value of the average thickness of the keratin bar in the sample, 2. Average soft block thickness – a value of the average thickness of the air bubble (in sphere type nanostructure) or air channel (in channel type nanostructure) embedded in the keratin. 3. Long period – a distance between the midpoint of one keratin bar and the nearest neighbouring keratin bar. Long period is used to calculate average soft block thickness by subtracting average hard block thickness from itand to calculate filling fraction. 4. Filling fraction - is calculated by dividing average hard block thickness by long period. It is a value indicating the percent material in the region containing the nanostructure. All four of the variables extracted from the correlation analysis were averaged for each species (Supplement material: Table S3.). The representation of the 3-D nanostructure and visual depiction of the variables is represented in the Fig. 2.

**Figure 2.**
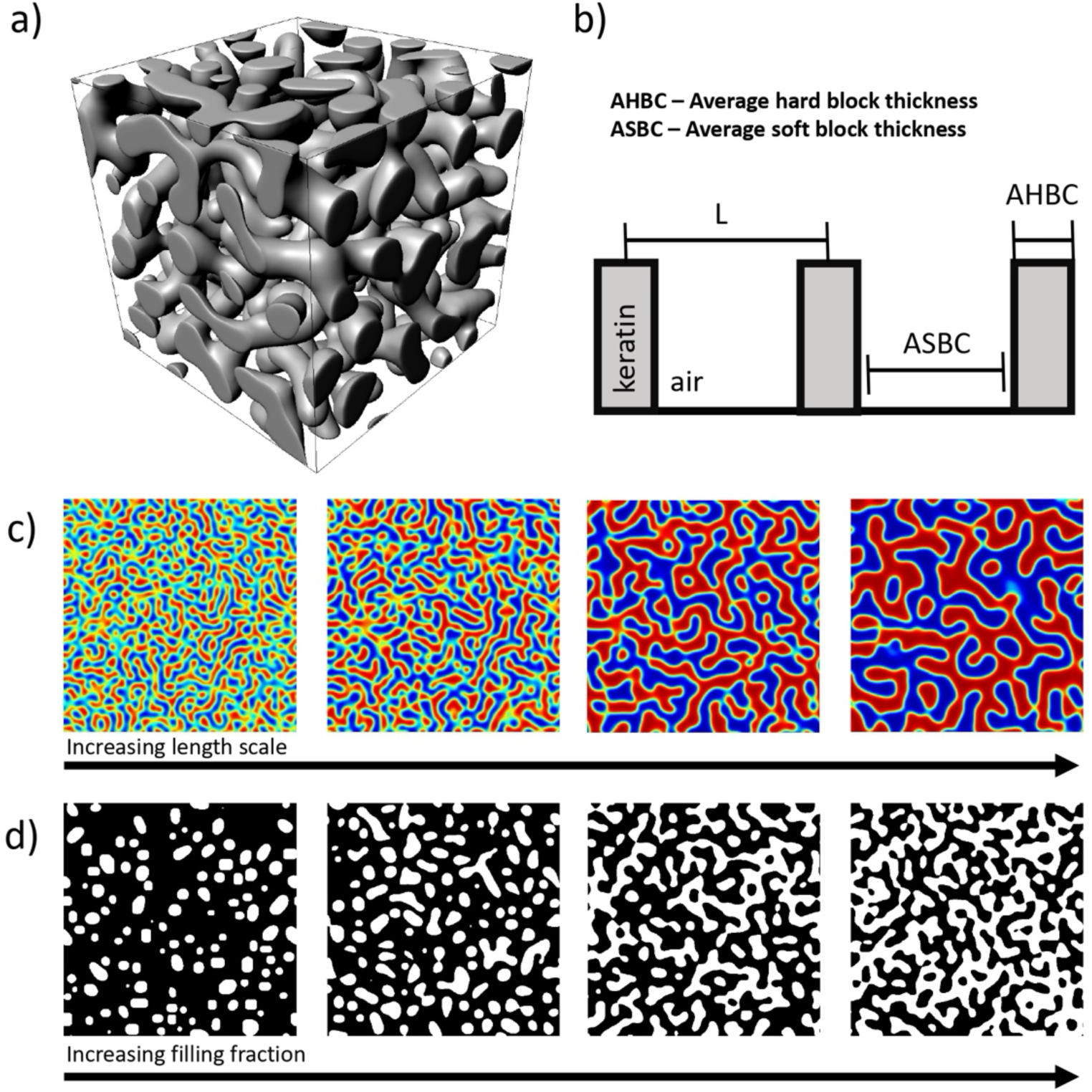
Visualization of the colour producing nanostructure and the variables extracted from the one-dimensional correlation analysis that describe its properties. Panel (a) shows a render of the channel-type nanostructure involved in the production of the colour blue. Keratin is shaded grey and unshaded area represents air. Panel (b) shows a 2-D representation of the 3-D keratin air and channel nanostructure. On the image, L stands for the long period, i.e. length between two keratin bars; ASBC is an average hard block thickness (keratin); ASBC is an average soft block thickness (air). Panel (c) is a representation of the filling fraction variable where red is the keratin and blue is the air.The length scales of the elements of the nanostructure do not change across the panels, but the percentage of the material filling the observed area does. Panel (d) is a representation of the increase in the length scale of the elements of the nanostructure. Black areas are keratin and white areas are air. Across the panels, an average length scale of these elements is increasing.

### Phylogenetic Generalized Least Squares (PGLS)

We used Phylogenetic Generalized Least Squares (PGLS) for the three analyses described below (Grafen & Hamilton, 1989) as implemented in the R package caper (Orme et al., 2013). In all cases we used molecular phylogenies of Tanagers available from birdtree.org (Jetz et al., 2012), as a phylogenetic framework. We downloaded 1000 random trees and extracted the maximum clade credibility tree in R using the maxCladeCred function from the phangorn package (Schliep, 2011).

In the first analysis to test which variables predict colour variation across blue, slate and grey colour, we used a multipredictor model with PC1 (approximating chromatic variation of the feathers, i.e. hue and saturation) of all three colours as a response variable and variables approximating nanostructure as a predictor variable (nanostructure complexity, average soft block thickness, average hard block thickness, filling fraction, and I_o_ (first scattering feature), summarized in Table 1. Since PC1 represents measurement of chromatic variation across all colour categories, with this analysis we will investigate which variables approximating nanostructure are important for the evolution of grey – slate – blue transition.

Next, we used a multipredictor model in PGLS to test which elements of the nanostructure influences variation in the chromatic component of the colour within blue (second analysis) andslate colour category (third analysis) separately. For this analysis, we used variables approximating nanostructure as a predictor variable (nanostructure, average soft block thickness, average hard block thickness, filling fraction and I_o_), and PC1 of a specific colour category as a response variable (i.e. PC1 of only blue colour and PC1 of only slate colour). With this analysis we wanted to explore what variables are affecting variation in individual colour and therefore are important for the evolution of hue and saturation (as approximated by PC1) within each colour category.

## Results

### Grey – slate – blue colour space

The first two principal components explained 97.5% of the variance in the raw cone-catch values: u, s, m, l of the measured feathers (Supplement material: Table S2; Fig. 3, a-c) with PC1 explaining 79.1 % and PC2 explaining 18.2% of the variance respectively (Supplement material: Table S4). Raw cone-catchvalues are obtained by transforming reflectance data measured by spectrometer (as outlined in the section Reflectance data). Since PC1 explained a high percentage of the variance in the raw cone-catch value data, we decided to use PC1 as a variable explaining chromatic variation of colour in further analysis. PC1 is one variable representing both hue and saturation (chromatic variation) of a certain feather. Lower values of PC1 indicated greater stimulation of s and u cones (blue and UV colouration), while higher values of PC1 indicated greater stimulation of m and l cones (red andgreen colouration). PC1 therefore aligns well with a grey – slate – blue transition with grey colour data associated with the highest PC1 values, slate colour data in the middle, and blue colour associated with the lowest PC1 values (Fig. 3, d).

**Figure 3.**
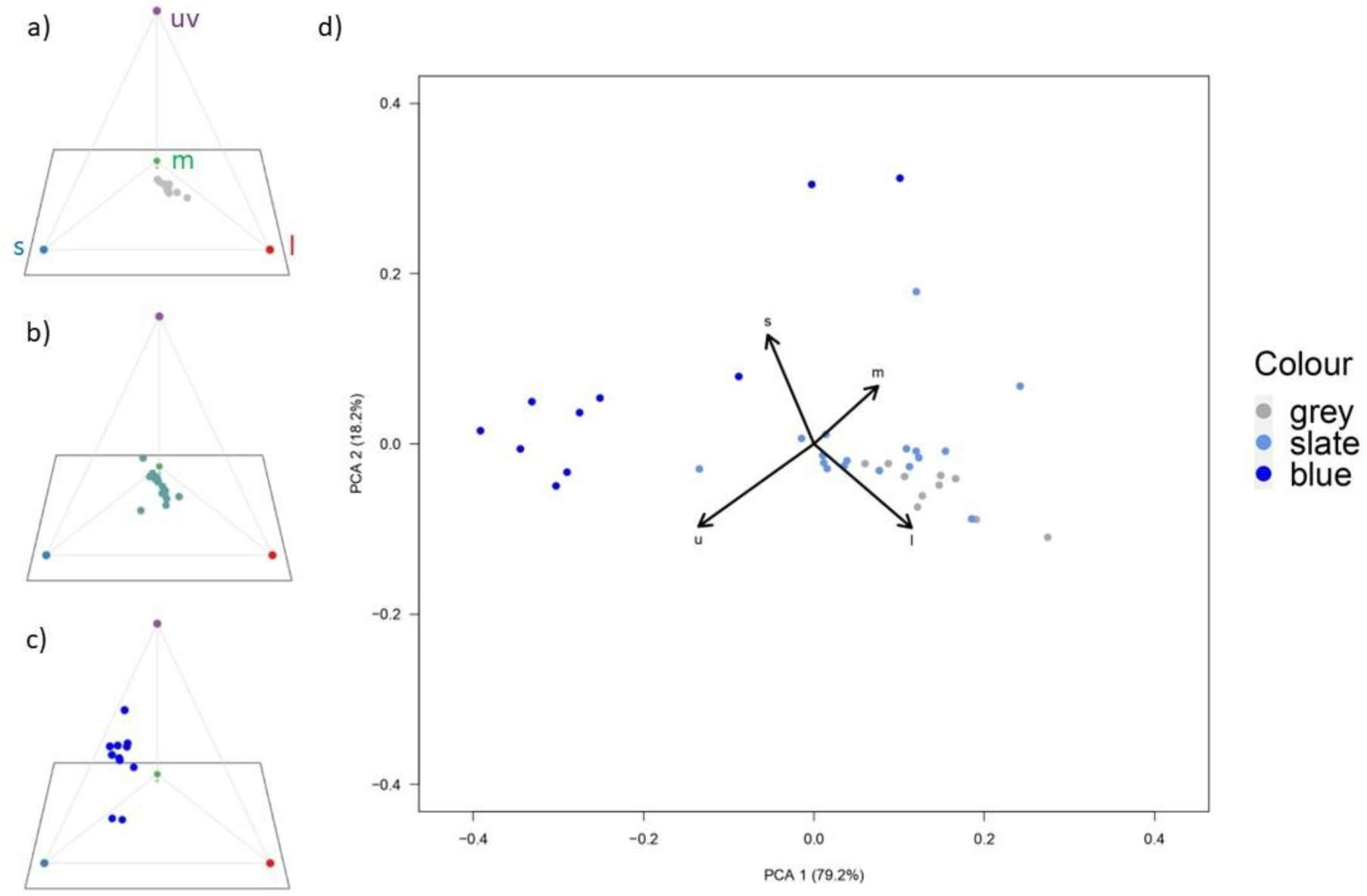
Panels a-c show the datapoints in avian tetrahedral colourspace for grey (a), slate (b) and blue (c) colour. The cone catch values describe every point in these 3 panels (u, s, m and l). Panel d shows principal components (PC) of cone catch values for all the feathers across all species. Each point in the plot represents one of the 38 feather samples measurements with point colour indicating which colour category a measurement belongs to (blue, slate or grey). PC1 explains the variation of colour scores. A higher PC1 value indicates a tendency toward m and l cone stimulations (grey colour in our case), while lower PC1 scores indicate a tendency towards blue and UV colour (blue in our case). Slate colour data points are roughly positioned between the data points for blue and grey colours.

### Description of nanostructural elements of feathers

We analysed all scattering profiles with the python code to detect peaks and shoulders. We divided the scattering profiles into categories according to the level of nanostructure detected and named that variable nanostructure complexity. The nanostructure complexity ranges from 0 (nanostructure is not detected) to 13 (nanostructure with the highest order features is present). Scores of 4, 7, 9 and 12 are not possible. The entire list of feathers and their scoring systems is in Additional table 4. (https://figshare.com/s/1110fce894e65a69c329), while a summary is presented in Fig. 4 and Supplement material: Table 1.

**Figure 4.**
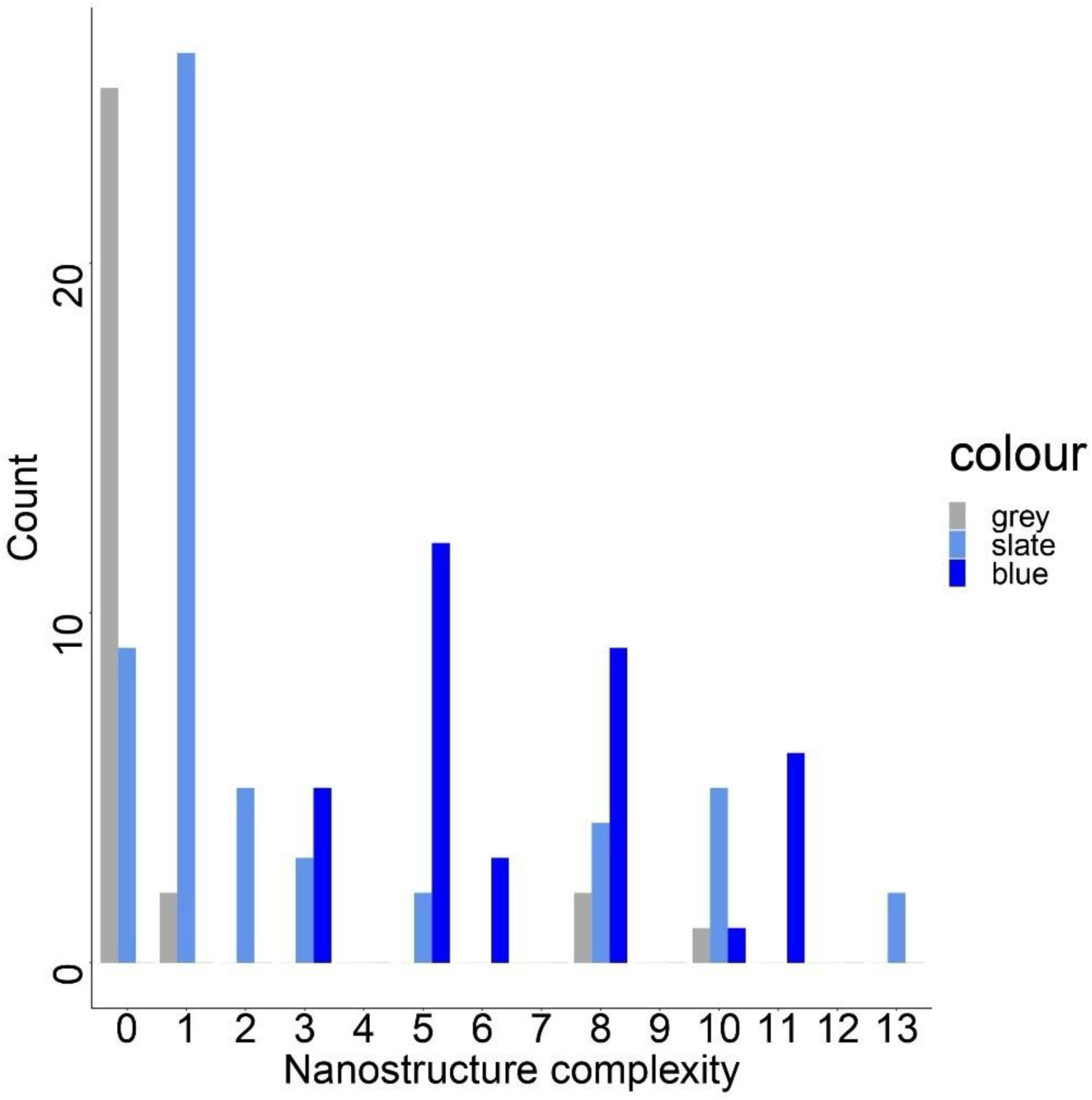
Histogram of the number of feathers (y axis) detected across all the feather samples for each category of nanostructure complexity variable (x axis). In short, every species was sampled with3 feathers, and we analysed 3 frames per each feather, making 117 feathers in total with 351 frames. Here, a feather was counted in certain nanostructure complexity category if at least one of the frames was detected belonging to that category. Feathers that did not have all three frames belonging to a same category are: *Chlorophanes spiza* (605), *Anisognathus igiventris* (608), *Pipraeidea melanoto* (612, 614), *Thraupis episcopus* (538), *Diglossa sittoides* (574), *Diglossa caerulescens* (577, 579), *Conirostrum cinerum* (590). In the brackets, a feather number as indicatedin the Additional table 4 (https://figshare.com/s/1110fce894e65a69c329).

### Phylogenetic generalised least square (PGLS) analysis results

The overview of the results is presented in the Fig. 5. Fig. 6-7 represent the effects of variables that showed significant correlation with colour variables. The full details of the analysis (*p*-values, parameter estimates and *R*^2^ values) are reported in the Supplement material: Table S4.

**Figure 5.**
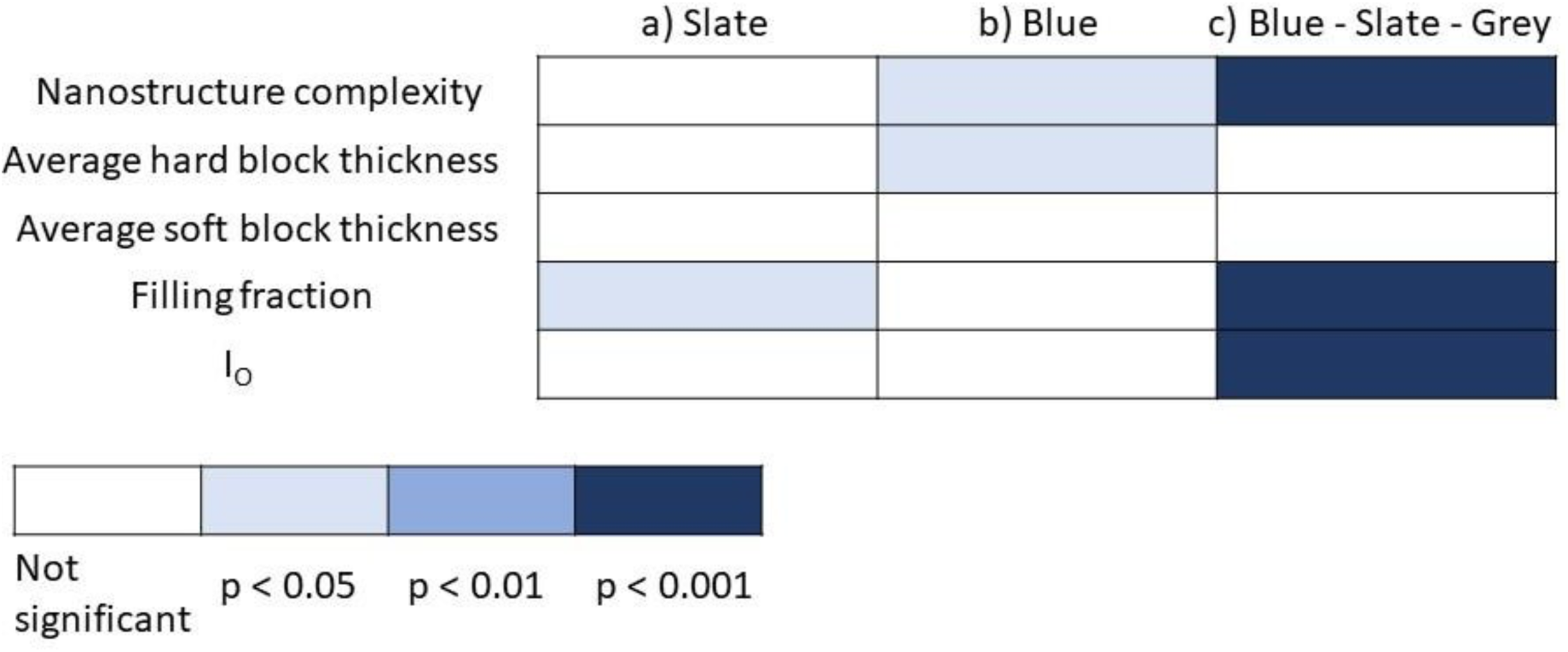
Multipredictor model results summary. All three panels represent values of PC1, with the panel a representing value only for slate colour, panel b only for the blue colour, and panel c representing combined values for grey, slate, and blue colour. Predictor variables are represented as rows with their names indicated further left. The colour of the squares represents the significance of the results, as indicated by the figure legend in the bottom left corner.

**Figure 6.**
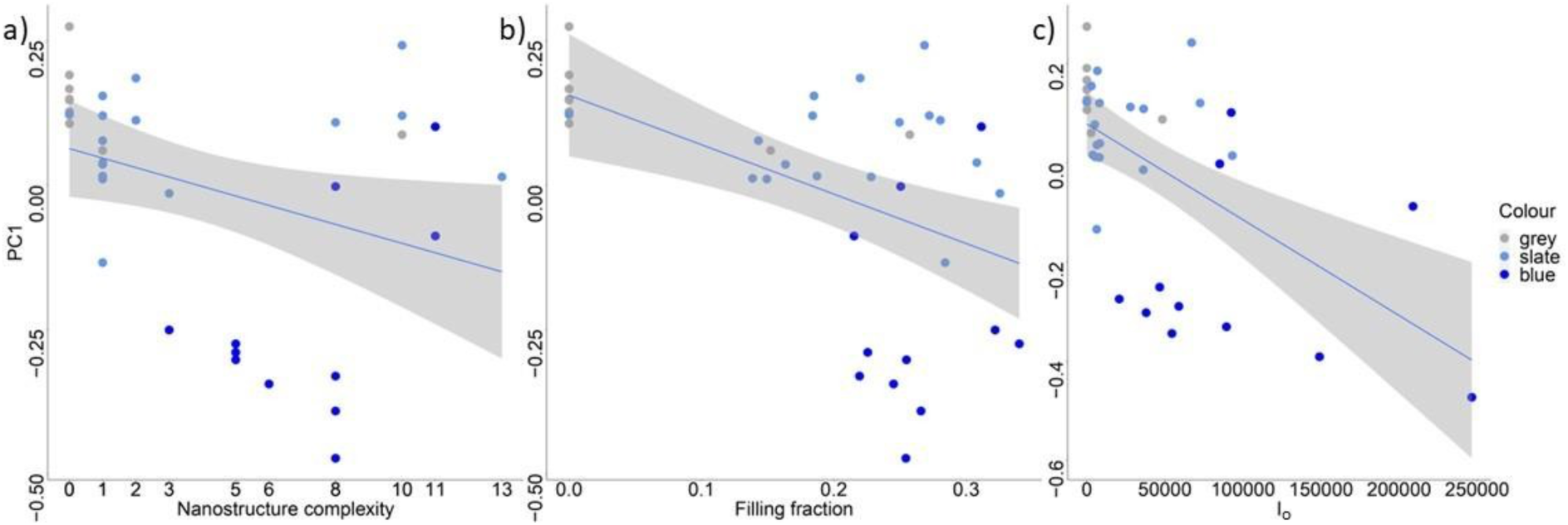
Predictors of PC1 for blue-slate-grey colour variation: a) nanostructure complexity, b) filling fraction, and c) I_o_. Within each panel, each point represents a species, and the colour of each point represents the colour category a measurement belongs to.

**Figure 7.**
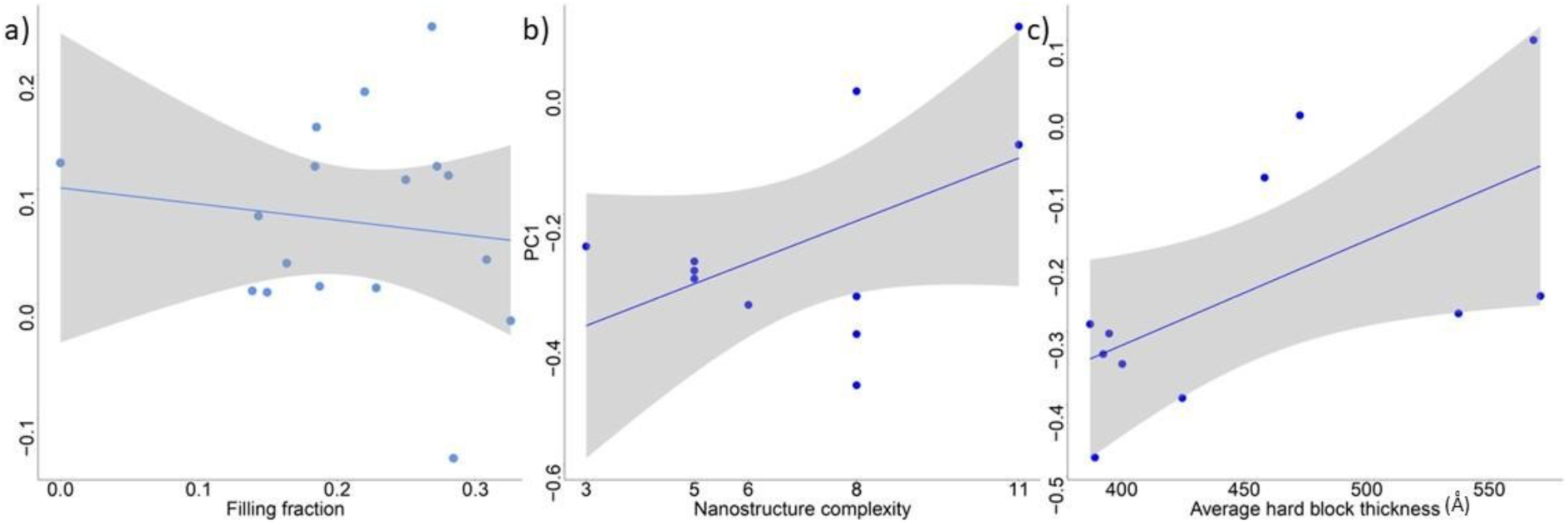
Predictors of PC1 for slate colour (a) and blue colour (b, c). Within each panel, each point represents a species. The predictor of slate colour PC1 variation is filling fraction (a), while predictors for PC1 of blue colour are nanostructure complexity (b) and average hard block thickness (c).

In the first analysis (Fig. 5, c), we used multipredictor PGLS analysis to assess which feather nanostructure variables correlated with the variation in the chromatic component of colour between colour categories as approximated by PC1. Nanostructure complexity (p = 0.0008953; slope = 2.9624e-02 (+/- 7.9681e-03)), filling fraction (p = 4.45E-08; slope = −2.9664e+00 (+/- 4.8490e-01)) and I_o_ (p= 0.0005619; slope = −1.9381e-06 (+/- 5.0592e-07)) showed significant association with the variation of the PC1 variable (Fig. 6, a–c). PC1 declines with increasing nanostructure complexity, filling fraction, and I_0_.

In the following analysis, we analysed slate and blue colour separately (i.e. in the analysis of slate colour, we analysed PC1 for only slate colour and in the analysis of blue colour, we analysed PC1 values for only blue colour as a response variable) (Fig. 5, a-b). For the slate colour analysis (Fig. 5, a; Fig. 7, a), only filling fraction (p = 0.01399, slope = −1.3408e+00 (+/- 4.8115e-01)) had a significant relationship with variation in PC1 (Fig. 7, a; Supplement material: Table S4, a). For a decrease in the value of PC1, there was an increase in the filling fraction value. For the blue colour analysis (Fig. 5, b), nanostructure complexity (p = 0.02315; slope = 4.0498e-02 (+/- 1.6217e-02)) and average hard block thickness (p = 0.01042; slope = 4.7362e-03 (+/- 3.3676e-03)) had a significant association with variation in PC1 (Fig. 7, b – c; Supplement material: Table S4, b). For an increase in the value of PC1, an increase in values of nanostructure complexity and average hard block thickness was detected.

Overall, explanatory power (R^2^) was greatest for model explaining variation in blue colour, followed by all three colours combined (blue-slate-grey) and finally model involving only slate colour had the lowest explanatory power (Supplement material: Table S4).

## 4.5. Discussion

We analysed the spongy structure of medullary keratinocytes in feather barbs from three broad colour groups (blue, slate and grey) to assess the mechanisms underpinning colour evolution from pigmentary grey to structural blue as well as variation within colour classes along this continuum. To do this we first quantified the absence or presence of nanostructure and classified the level of nanostructure present. We then quantified length scales and properties of the colour producing nanostructure, i.e. average hard block thickness, average soft block thickness, filling fraction and I_o_.

Correlates of variation in chromatic component of colour encompassing all three colour categories included nanostructure complexity, filling fraction and I_o_, while average hard block thickness and average soft block thickness showed no significant association. This indicates that it is the ratio of keratin to air that is more important than variation in keratin thickness for colour variation. However, patterns across the colours do not translate to within-colour categories correlates, i.e. those for blue and slate colour individually. PC1 values for blue colour were correlated with nanostructure level and average hard block thickness, while slate colour PC1 showed correlationwith the filling fraction. This pattern shows that while multiple components of variability in medullary cells spongy layer are needed for evolutionary transitions between blue, slate and grey occur, a more limited number of variables account for the variation in chromatic component of colour within the colour categories themselves.

Evolutionary transitions from pigmentary to structural colour have previously been detected in birds’ plumage (Shawkey et al., 2006; Driskell et al., 2010; Doucet et al., 2004). Our results indicate that for the transition from pigmentary grey towards structural blue colour, multiple variables describing spongy layer are important. I_o_ (thickness of the spongy layer), filling fraction and degree of order (nanostructure complexity) all increase as colour tends towards blue (PC1 decreases). Separately, for both blue and slate colour, the I_o_ (thickness of the spongy layer) does not show a correlation with PC1. This could indicate that there might be a critical length scale of the nanostructure that is important for the evolutionary transition from grey to blue to happen. Increasing thickness of the spongy layer (correlated with the increase in I_o_) will result in greater reflectance across the short-wavelength range, i.e. blue and UV (Fan et al., 2019). Filling fraction is a measure of what volume fraction is occupied or filled by the biopolymer (keratin). To produce white colour in some species of beetles, it has been proposed that a filling fraction of 31 – 34 % is responsible for the colour production, while simulated results indicate a theoretical maximum reflectance from a spongy nanostructure at 25% (Burg et al., 2019). This is observed in our results as well, i.e. increase in filling fraction from 0 (for *Sporophila caerulescens* grey feather) through 0.1386 (13.86% for *Catemina analis* slate rump feather) to 0.34012 (34.012% for *Diglossa cayana* blue rump feather) is observed with decreasing PC1 (moving towards blue colour in the colourspace). This results further confirms nanostructural resemblance in spongy structure between blue and white colour in bird’s feathers as previously observed in amelanotic Steller’s jay (*Cyanocitta stelleri*) and in swallow tanager (*Tersina viridis*) (Bazzano et al., 2021; Shawkey & Hill, 2006). In both cases, white and blue feathers have similar peak in reflectance in blue part of the spectrum, but the pronunciation of the peak in blue feathers is due to the underlying melanin layer which is lacking in white feathers. Finally, the value of nanostructural complexity showed an increase with decreasing PC1 values, and this could indicate that blue colour is associated with structural uniformity and increased order of the nanostructures. Overall, changes in many variables explaining spongy barb nanostructure have proven to be important for the evolution of grey-slate-blue continuum in the colourspace.

Previous research into changes in nanostructural parameters between different hues of non-iridescent structural colour revealed that variation in many nanostructural elements, rather than a change in single parameter, is responsible for observed colour diversity (Fan et al., 2019). These parameters involve the thickness of the outer layer of keratin (above colour producing nanostructure), spatial frequency and thickness of the keratin and air matrix, as well as the amount of melanin beneath the colour producing nanostructure. Our results are focused only on the blue colour and show that two main components for colour production are nanostructure complexity andhard block thickness (Fig. 5, b; Fig. 7, b-c). The increase in PC1 follows increasing hard block thickness indicating that thicker keratin bars in either channel or sphere type spongy layer would shift away from blue and UV cone stimulations. Increases in the level of nanostructure also followthe same trend. Surprisingly, we did not find a thickness of the spongy layer as a correlate of PC1 of the colour blue as opposed to the previous research (Fan et al., 2019). This could be explained by theabsence of other structural colours from our dataset, namely purple. Thicker spongy layer would increase reflectance in the short wavelengths (Fan et al., 2019), meaning that the spongy structure length scale could be correlated if we had a broader range of structural colours within our dataset. Nevertheless, this variable proved to be important for the transition into blue colour from slate (as showed by our results).

In previous research on the nanostructure of slate colour it has been identified that this colour category is characterised by more rudimentary and highly disordered versions of the channel and sphere type nanostructures that are found in the blue feathers (Saranathan et al., 2012). Nevertheless, it seems that these feathers still have nanostructure ordered enough to produce colour by coherent scattering. The only variable that correlates with PC1 for slate colour is filling fraction where higher values of filling fraction are associated with lower values of PC1 for slate colour. Within slate colour category, higher values of filling fraction correlate with the lower PC1 values showing more inclinations toward colour blue (i.e. blue and UV cone stimulation). As explained previously, filling fraction is the value that indicates the filling of the volume of thecrystalline structure with its constituent elements, i.e. keratin in our case. Increasing filling fraction has been shown to be important in evolution of colour blue (this research) while it is not important for a hue variation within blue colour category. A limitation of our research is not knowing the location of melanosomes within the feathers. It is known from literature that coherent scattering that produces blue colour can be masked by melanin deposition and in that case the feather is black (Doucet et al., 2004; Driskell et al., 2010). Whether this rudimentary spongy layer detected in the slate feathers has a melanosome deposition above it that participate in the colour production by interfering with the colour produced from the spongy layer is yet to be seen. Nevertheless, the fact that variation in PC1 for slate colour correlates with filling fraction indicates that the spongy layer is ordered enough to participate in colour production (giving the slight blue of the slate colour).

Our results suggest that the parameters of spongy structure that influence colour variation between colour categories (blue-slate-grey) differ from parameters that influence colour variation within colour categories (blue and slate). Structural colours are intrinsically linked to their underlying nanostructure (Prum, 2006). It has been shown that small changes in their nanostructures will leadto a change in the colour produced and, therefore, the signal emitted in the environment (Fan et al., 2019; Saranathan et al., 2012). Development of the spongy structure is proceeding without active cellular processes, i.e. by phase separation of the mixture of keratin and air in the medullary cells (Prum et al., 2009). These self-guided processes could theoretically lead to complete unmixing of thesolution and loss of nanostructure arrangement necessary for colour production (Jones, 2002; Prum et al., 2009). It is still debated what causes halts in the phase separation during feather growth (and colour production), but it is known that these physical processes are deterministic, and there is little opportunity for a variation in the outcome of the development once the process is initiated (Jones, 2002; Prum et al., 2009). Our results indicate that the inherent issue with the phase separation (its deterministic nature) and control over the variation in hue within and between colour categories could be overcome by varying different elements (slate colour results) or combinations of elements (blue-slate-grey and blue colour results) of spongy structure in medullary cells. The variation in multiple elements of the nanostructure rather than binary presence/absence scheme for productions of different hues has been already confirmed for non-iridescent structural colours (Fan et al., 2019). It seems that similar processes are involved in their evolution and here we propose thatthis is a natural consequence of utilizing basic physical processes during feather development.

