## Supplementary material for "The mechanistic basis of evolutionary transitions between grey, slate, and blue colour in Tanagers (Thraupidae)": The mechanistic basis of evolutionary transitions between grey, slate, and blue colour in Tanagers (Thraupidae)_supplement_material.docx

### Evolutionary dynamics of pigmentary grey and non- iridescent structural blue colouration in Tanagers (family: Thraupidae) – Supplement material

**Table S1.** Sampling details of feathers from museum skins in the the Zoological Museum, Natural History Museum of Denmark, University of Copenhagen. Every row represents on feather and columns indicate details about the specimen (Species, Sex, Storage number, Tag details) and feather samples (Colour, Body part).

| Species | Sex | Body part | Storage number | Tag details | Colour |
| --- | --- | --- | --- | --- | --- |
| Tangara vitriolina | Unsexed | Throat | NA | 8.4.1913.220 | Slate |
| Tangara vitriolina | Unsexed | Throat | NA | 8.4.1913.219 | Slate |
| Tangara vitriolina | Unsexed | Throat | NA | 8.4.1913.221 | Slate |
| Catemina inornata | Male | Mantle | 105120 | 6.5.2010 | Slate |
| Catemina inornata | Male | Mantle | 105117 | 6.5.2010 | Slate |
| Catemina inornata | Male | Mantle | 105118 | 6.5.2010 | Slate |
| Poospiza cinera | Unsexed | Mantle | 20.578 | NA | Slate |
| Poospiza cinera | Male | Mantle | 20.576 | NA | Slate |
| Poospiza cinera | Unsexed | Mantle | 20.580 | NA | Slate |
| Diglossa cyanea | Male | Rump | 104778 | 20.11.1984.569 | Blue |
| Diglossa cyanea | Male | Rump | 104781 | 22.4.2010 | Blue |
| Diglossa cyanea | Male | Rump | 104782 | 22.4.2010 | Blue |
| Xenodacnis parina | Male | Mantle | 103020 | 20.3.2009-9 | Blue |
| Xenodacnis parina | Male | Rump | 103022 | 20.3.2009-11 | Blue |
| Xenodacnis parina | Male | Rump | 103021 | 20.3.2009-10 | Blue |
| Sporophila caerulescens | Male | Mantle | 20.468 | 26.4.1937.692 | Grey |
| Sporophila caerulescens | Male | Mantle | 20.467 | 26.4.1937.691 | Grey |
| Sporophila caerulescens | Male | Mantle | 20.462 | 26.4.1937.686 | Grey |
| Phygilus unicolor | Male | Mantle | 64080 | 26.8.1983-167 | Grey |
| Phygilus unicolor | Male | Mantle | 105109 | 6.5.2010 | Grey |
| Phygilus unicolor | Male | Mantle | 64078 | 26.8.1983-165 | Grey |
| Coryphospingus plieatus | Male | Rump | 9796 | NA | Grey |
| Coryphospingus plieatus | Male | Rump | 9792 | NA | Grey |
| Coryphospingus plieatus | Male | Rump | 9797 | NA | Grey |
| Diuca diuca | Male | Breast | 9720 | NA | Grey |

| Diuca diuca | Male | Breast | 9721 | 28.6.1937.7 | Grey |
| --- | --- | --- | --- | --- | --- |
| Diuca diuca | Male | Breast | 9717 | NA | Grey |
| Poospiza hypochondria | Unsexed | Mantle | 104995 | 3.5.2010 | Grey |
| Poospiza hypochondria | Female | Mantle | 104996 | 3.5.2010 | Grey |
| Poospiza hypochondria | Male | Mantle | 104998 | 3.5.2010 | Grey |
| Saltator albicollis | Unsexed | Nape | 20.274 | NA | Grey |
| Saltator albicollis | Unsexed | Nape | 20.260 | NA | Grey |
| Saltator albicollis | Male | Nape | 20.278 | NA | Grey |
| Neothraupis fasciata | Male | Mantle | NA | NA | Grey |
| Neothraupis fasciata | Male | Mantle | NA | NA | Grey |
| Neothraupis fasciata | Male | Mantle | NA | NA | Grey |
| Sporophila plumbea | Male | Mantle | 20.374 | NA | Grey |
| Sporophila plumbea | Male | Mantle | 20.370 | NA | Grey |
| Sporophila plumbea | Male | Mantle | 20.373 | NA | Grey |
| Thraupis sayaca | Male | Throat | NA | 26.4.1937.578 | Grey |
| Thraupis sayaca | Male | Throat | NA | 26.4.1937.579 | Grey |
| Thraupis sayaca | Male | Throat | NA | 26.4.1937.577 | Grey |
| Phryglius alaudinus | Unsexed | Throat | 9765 | 15.2.1915.211 | Slate |
| Phryglius alaudinus | Male | Throat | 67932 | 27.3.1979.138 | Slate |
| Phryglius alaudinus | Male | Throat | 9761 | NA | Slate |
| Thraupis cyanocephala | Male | Breast | NA | 30.1.91-41 | Slate |
| Thraupis cyanocephala | Male | Breast | 103009 | 20.9.1980.48 | Slate |
| Thraupis cyanocephala | Male | Breast | NA | 30.1.91-40 | Slate |
| Thraupis episcopus | Male | Throat | NA | 30.1.91-39 | Slate |
| Thraupis episcopus | Male | Throat | NA | 20.9.1980.46 | Slate |
| Thraupis episcopus | Male | Mantle | NA | 30.1.91-38 | Slate |
| Tangara heinei | Male | Breast | NA | NA | Slate |
| Tangara heinei | Male | Breast | NA | 30.1.1991-287 | Slate |
| Tangara heinei | Male | Breast | NA | 15.2.1915.185 | Slate |
| Catamblyrhynchus diadema | Male | Mantle | 104900 | 28.4.2010 | Slate |
| Catamblyrhynchus diadema | Unsexed | Mantle | 9647 | 8.2.1921.490 | Slate |
| Catamblyrhynchus diadema | Unsexed | Mantle | 64085 | 26.8.1983-156 | Slate |
| Catamenia analis | Male | Rump | 105122 | 6.5.2010 | Slate |
| Catamenia analis | Male | Rump | 105114 | 6.5.2010 | Slate |
| Catamenia analis | Male | Rump | 10515 | 6.5.2010 | Slate |
| Diglossa lafresnayi | Unsexed | Wing covert | NA | 1.6.1930.366 | Slate |
| Diglossa lafresnayi | Unsexed | Wing covert | NA | NA | Slate |
| Diglossa lafresnayi | Unsexed | Wing covert | NA | 8.2.1921.349 | Slate |
| Diglossa albilatera | Unsexed | Breast | NA | NA | Slate |
| Diglossa albilatera | Male | Breast | 104797 | 22.4.2010 | Slate |
| Diglossa albilatera | Male | Breast | NA | NA | Slate |
| Diglossa brunneiventris | Male | Rump | 104899 | 27.4.2010 | Slate |
| Diglossa brunneiventris | Male | Rump | 103034 | 20.3.2009-21 | Slate |
| Diglossa brunneiventris | Male | Rump | 103040 | NA | Slate |
| Diglossa sittoides | Male | Nape | NA | 8.2.1921.353 | Slate |
| Diglossa sittoides | Male | Nape | NA | NA | Slate |

| Diglossa sittoides | Male | Mantle | NA | NA | Slate |
| --- | --- | --- | --- | --- | --- |
| Diglossa caerulescens | Male | Breast | 104791 | 22.4.2010 | Slate |
| Diglossa caerulescens | Male | Breast | 104793 | 22.4.2010 | Slate |
| Diglossa caerulescens | Male | Breast | NA | NA | Slate |
| Oreomanes fraseri | Male | Nape | 104831 | 23.4.2010 | Slate |
| Oreomanes fraseri | Male | Nape | 104822 | 23.4.2010 | Slate |
| Oreomanes fraseri | Male | Nape | 104839 | 26.4.2010 | Slate |
| Conirostrum cinerum | Male | Nape | 104802 | 27.3.1979.130 | Slate |
| Conirostrum cinerum | Male | Nape | 104801 | 23.4.2010 | Slate |
| Conirostrum cinerum | Male | Nape | 104803 | 23.4.2010 | Slate |
| Conirostrum sitticolor | Unsexed | Rump | 104804 | 23-4-1985-18 | Blue |
| Conirostrum sitticolor | Male | Rump | 103028 | 20.3.2009-17 | Blue |
| Conirostrum sitticolor | Male | Rump | 104805 | 23.4.2010 | Blue |
| Dacnis cayana | Male | Breast | 137319 | 8.6.2011.20 | Blue |
| Dacnis cayana | Male | Breast | NA | 27.3.1979.131 | Blue |
| Dacnis cayana | Male | Breast | NA | 20.9.1980.38 | Blue |
| Cyanerpes cyaneus | Male | Breast | 59979 | 1.7.1971.579 | Blue |
| Cyanerpes cyaneus | Male | Breast | NA | 25.12.1922.99 | Blue |
| Cyanerpes cyaneus | Male | Breast | 59980 | 1.7.1971.580 | Blue |
| Chlorophanes spiza | Male | Breast | NA | NA | Blue |
| Chlorophanes spiza | Male | Breast | 103018 | NA | Blue |
| Chlorophanes spiza | Male | Breast | NA | 1.6.1930.373 | Blue |
| Anisognathus igniventris | Male | Rump | 102988 | 19.3.2009-2 | Blue |
| Anisognathus igniventris | Male | Rump | NA | 20.1.1982 | Blue |
| Anisognathus igniventris | Male | Rump | 104882 | 26.4.2010 | Blue |
| Pipraeidea melanota | Male | Rump | NA | NA | Blue |
| Pipraeidea melanota | Male | Rump | NA | NA | Blue |
| Pipraeidea melanota | Unsexed | Rump | NA | NA | Blue |
| Tangara chilensis | Male | Throat | NA | 1.6.1930.393 | Blue |
| Tangara chilensis | Unsexed | Throat | NA | 1.6.1930.396 | Blue |
| Tangara chilensis | Unsexed | Throat | NA | 1.6.1930.395 | Blue |
| Tangara nigroviridis | Unsexed | Throat | NA | 8.4.1913.203 | Blue |
| Tangara nigroviridis | Male | Throat | 92.806 | 18.12.1998-16 | Blue |
| Tangara nigroviridis | Male | Throat | NA | 30.1.91-137 | Blue |
| Poospiza schistacea | Unsexed | Mantle | 20.580 | NA | Slate |
| Poospiza schistacea | Unsexed | Mantle | 20.578 | NA | Slate |
| Poospiza schistacea | Male | Mantle | 20.576 | NA | Slate |
| Paroaria coronata | Male | Mantle | 9311 | NA | Grey |
| Paroaria coronata | Male | Mantle | 9310 | 2.4.1917.1 | Grey |
| Paroaria coronata | Male | Mantle | 9303 | NA | Grey |
| Thraupis bonariensis | Male | Throat | NA | R 2-8-1907 | Blue |
| Thraupis bonariensis | Male | Throat | 103013 | 20.3.2009-1 | Blue |
| Thraupis bonariensis | Male | Throat | 92.805 | 18.1.1998-15 | Blue |
| Charitospiza eucoma | Male | Mantle | 9772 | NA | Grey |
| Charitospiza eucoma | Male | Mantle | 9773 | NA | Grey |
| Charitospiza eucoma | Male | Mantle | 9774 | NA | Grey |

**Table S2.** Colour measurements for feather samples in Tanagers. Each row represents average colour measurement per species from 3 feathers. First column shows Latin name of the species, second patch where the feathers are sampled from, third colour of the feathers, columns 4 – 7 show cone catch values, eight column shows Projection values, column 9 – 10 show PC1 and PC2 from Principal component analysis.

| Latin name | Patch | Colour | u | s | m | l | Projection_va  lues | PC1 | PC2 |
| --- | --- | --- | --- | --- | --- | --- | --- | --- | --- |
| Diglossa cyanea | rump | blue | 0.386 | 0.405 | 0.151 | 0.059 | 0.191 | -0.275 | 0.037 |
| Xenodacnis parina | mantle /  rump | blue | 0.366 | 0.398 | 0.180 | 0.055 | 0.203 | -0.251 | 0.054 |
| Conirostrum sitticolor | rump | blue | 0.455 | 0.339 | 0.136 | 0.070 | 0.322 | -0.303 | -0.049 |
| Dacnis cayana | breast | blue | 0.276 | 0.301 | 0.343 | 0.080 | 0.397 | -0.088 | 0.079 |
| Cyanerpes cyaneus | breast | blue | 0.465 | 0.445 | 0.070 | 0.021 | 0.110 | -0.392 | 0.015 |
| Chlorophanes spiza | breast | blue | 0.010 | 0.453 | 0.420 | 0.118 | 0.095 | 0.101 | 0.312 |
| Anisognathus igniventris | rump | blue | 0.454 | 0.396 | 0.111 | 0.040 | 0.208 | -0.345 | -0.006 |
| Pipraeidea melanonota | rump | blue | 0.412 | 0.440 | 0.117 | 0.030 | 0.119 | -0.331 | 0.050 |
| Tangara chilensis | throat | blue | 0.626 | 0.309 | 0.047 | 0.018 | 0.382 | -0.473 | -0.156 |
| Tangara nigroviridis | throat | blue | 0.060 | 0.526 | 0.311 | 0.103 | 0.052 | -0.003 | 0.305 |
| Thraupis bonariensis | crown | blue | 0.437 | 0.348 | 0.144 | 0.071 | 0.305 | -0.290 | -0.033 |
| Sporophila caerulescens | mantle | grey | 0.136 | 0.240 | 0.289 | 0.335 | 0.521 | 0.149 | -0.037 |
| Phrygilus unicolor | mantle | grey | 0.192 | 0.268 | 0.266 | 0.274 | 0.464 | 0.060 | -0.023 |
| Coryphospingus pileatus | rump | grey | 0.172 | 0.225 | 0.264 | 0.339 | 0.551 | 0.122 | -0.074 |
| Diuca diuca | breast | grey | 0.124 | 0.238 | 0.286 | 0.352 | 0.523 | 0.166 | -0.041 |
| Poospiza hypochondria | mantle | grey | 0.130 | 0.202 | 0.278 | 0.390 | 0.596 | 0.191 | -0.089 |
| Saltator albicollis | nape | grey | 0.089 | 0.153 | 0.319 | 0.438 | 0.693 | 0.275 | -0.110 |
| Neothraupis fasciata | mantle | grey | 0.160 | 0.234 | 0.267 | 0.338 | 0.531 | 0.127 | -0.061 |
| Sporophila plumbea | mantle | grey | 0.141 | 0.237 | 0.279 | 0.343 | 0.526 | 0.147 | -0.048 |
| Thraupis sayaca | throat | grey | 0.175 | 0.258 | 0.281 | 0.287 | 0.485 | 0.087 | -0.023 |
| Charitospiza eucosma | nape | grey | 0.167 | 0.247 | 0.278 | 0.309 | 0.507 | 0.106 | -0.038 |
| Catamenia inornata | mantle | slate | 0.145 | 0.256 | 0.291 | 0.307 | 0.488 | 0.123 | -0.016 |
| Poospiza cinerea | mantle | slate | 0.118 | 0.257 | 0.299 | 0.325 | 0.486 | 0.154 | -0.009 |
| Phrygilus alaudinus | throat | slate | 0.184 | 0.258 | 0.270 | 0.288 | 0.483 | 0.077 | -0.031 |
| Thraupis cyanocephala | breast | slate | 0.148 | 0.271 | 0.283 | 0.298 | 0.457 | 0.108 | -0.006 |
| Thraupis episcopus | throat/ma  ntle | slate | 0.225 | 0.262 | 0.315 | 0.198 | 0.476 | 0.014 | 0.011 |
| Tangara heinei | breast | slate | 0.048 | 0.392 | 0.342 | 0.218 | 0.216 | 0.120 | 0.179 |
| Catamblyrhynchus diadema | mantle | slate | 0.224 | 0.280 | 0.261 | 0.235 | 0.439 | 0.010 | -0.014 |
| Catamenia analis | rump | slate | 0.226 | 0.277 | 0.256 | 0.241 | 0.447 | 0.012 | -0.022 |
| Diglossa lafresnayii | wing_cove  rts | slate | 0.332 | 0.304 | 0.208 | 0.155 | 0.392 | -0.134 | -0.030 |
| Diglossa albilatera | breast | slate | 0.212 | 0.268 | 0.263 | 0.257 | 0.464 | 0.036 | -0.026 |
| Diglossa brunneiventris | rump | slate | 0.229 | 0.268 | 0.260 | 0.244 | 0.465 | 0.015 | -0.029 |
| Diglossa sittoides | nape /  mantle | slate | 0.206 | 0.271 | 0.266 | 0.257 | 0.457 | 0.039 | -0.020 |
| Diglossa caerulescens | breast | slate | 0.231 | 0.299 | 0.259 | 0.210 | 0.401 | -0.015 | 0.006 |
| Oreomanes fraseri | nape | slate | 0.135 | 0.200 | 0.281 | 0.384 | 0.599 | 0.185 | -0.088 |
| Conirostrum cinereum | nape | slate | 0.159 | 0.249 | 0.288 | 0.304 | 0.501 | 0.112 | -0.027 |

| Sporophila schistacea | mantle | slate | 0.143 | 0.263 | 0.291 | 0.303 | 0.474 | 0.120 | -0.009 |
| --- | --- | --- | --- | --- | --- | --- | --- | --- | --- |
| Tangara vitriolina | mantle | slate | 0.041 | 0.240 | 0.416 | 0.303 | 0.521 | 0.242 | 0.068 |

**Table S3.** Measurements of variables explaining properties of spongy layer for feathers in Tanagers. Every row represents an average value of the measurement for a species.

| Latin name | Patch | Colour | Nanostruct ure complexity | Long Period | Average Hard Block Thickne  ss | Average Soft Block Thickne  ss | Filling fraction | I_(max) | q_(max) |
| --- | --- | --- | --- | --- | --- | --- | --- | --- | --- |
| Diglossa_cyanea | rump | blue | 5 | 1580.86  0 | 537.452 | 1043.40  8 | 0.340 | 20779.73  9 | 0.003 |
| Xenodacnis_parina | mantle /  rump | blue | 3 | 1772.81  0 | 570.865 | 1201.94  5 | 0.322 | 46668.92  5 | 0.003 |
| Conirostrum_sitticolor | rump | blue | 5 | 1549.06  1 | 394.873 | 1154.18  8 | 0.255 | 37991.22  8 | 0.004 |
| Dacnis_cayana | breast | blue | 11 | 2129.78  3 | 458.322 | 1671.46  0 | 0.215 | 208921.3  55 | 0.003 |
| Cyanerpes_cyaneus | breast | blue | 8 | 1597.81  8 | 424.751 | 1173.06  7 | 0.266 | 149011.1  12 | 0.004 |
| Chlorophanes_spiza | breast | blue | 11 | 1823.65  0 | 568.041 | 1255.60  9 | 0.311 | 92476.24  1 | 0.003 |
| Anisognathus_igniventris | rump | blue | 6 | 1633.11  1 | 400.252 | 1232.85  9 | 0.245 | 54499.42  3 | 0.004 |
| Pipraeidea_melanonota | rump | blue | 8 | 1790.95  4 | 392.479 | 1398.47  4 | 0.219 | 89431.47  5 | 0.003 |
| Tangara_chilensis | throat | blue | 8 | 1528.86  6 | 389.079 | 1139.78  7 | 0.254 | 246532.3  22 | 0.004 |
| Tangara_nigroviridis | throat | blue | 8 | 1887.18  5 | 472.732 | 1414.45  2 | 0.251 | 85120.93  4 | 0.003 |
| Thraupis_bonariensis | crown | blue | 5 | 1715.94  3 | 387.121 | 1328.82  2 | 0.226 | 58920.09  6 | 0.003 |
| Sporophila_caerulescens | mantle | grey | 0 | 0.000 | 0.000 | 0.000 | 0.000 | 0.000 | 0.000 |
| Phrygilus_unicolor | mantle | grey | 1 | 4565.18  4 | 693.784 | 3871.40  1 | 0.152 | 2749.584 | 0.004 |
| Coryphospingus_pileatus | rump | grey | 0 | 0.000 | 0.000 | 0.000 | 0.000 | 0.000 | 0.000 |
| Diuca_diuca | breast | grey | 0 | 0.000 | 0.000 | 0.000 | 0.000 | 0.000 | 0.000 |
| Poospiza_hypochondria | mantle | grey | 0 | 0.000 | 0.000 | 0.000 | 0.000 | 0.000 | 0.000 |
| Saltator_albicollis | nape | grey | 0 | 0.000 | 0.000 | 0.000 | 0.000 | 0.000 | 0.000 |
| Neothraupis_fasciata | mantle | grey | 0 | 0.000 | 0.000 | 0.000 | 0.000 | 0.000 | 0.000 |
| Sporophila_plumbea | mantle | grey | 0 | 0.000 | 0.000 | 0.000 | 0.000 | 0.000 | 0.000 |
| Thraupis_sayaca | throat | grey | 10 | 2366.48  2 | 608.738 | 1757.74  4 | 0.257 | 48398.28  9 | 0.003 |
| Charitospiza_eucosma | nape | grey | 0 | 0.000 | 0.000 | 0.000 | 0.000 | 0.000 | 0.000 |
| Catamenia_inornata | mantle | slate | 0 | 0.000 | 0.000 | 0.000 | 0.000 | 0.000 | 0.000 |
| Poospiza_cinerea | mantle | slate | 1 | 4891.57  6 | 907.524 | 3984.05  2 | 0.185 | 2906.703 | 0.005 |
| Phrygilus_alaudinus | throat | slate | 1 | 4531.91  1 | 647.932 | 3883.97  9 | 0.143 | 5088.436 | 0.004 |
| Thraupis_cyanocephala | breast | slate | 8 | 1890.39  0 | 472.018 | 1418.37  3 | 0.250 | 36345.31  1 | 0.003 |
| Thraupis_episcopus | throat/man  tle | slate | 13 | 2245.88  8 | 512.694 | 1733.19  5 | 0.228 | 93194.20  9 | 0.003 |
| Tangara_heinei | breast | slate | 10 | 1920.84  3 | 522.688 | 1398.15  5 | 0.272 | 72541.88  3 | 0.003 |
| Catamblyrhynchus_diade  ma | mantle | slate | 1 | 4681.16  2 | 714.423 | 3966.73  9 | 0.149 | 8038.068 | 0.004 |
| Catamenia_analis | rump | slate | 1 | 4554.47  8 | 629.440 | 3925.03  7 | 0.139 | 5340.388 | 0.004 |

| Diglossa_lafresnayii | wing_cover  ts | slate | 1 | 1551.94  6 | 438.895 | 1113.05  1 | 0.284 | 6338.845 | 0.004 |
| --- | --- | --- | --- | --- | --- | --- | --- | --- | --- |
| Diglossa_albilatera | breast | slate | 1 | 4958.44  4 | 812.630 | 4145.81  4 | 0.164 | 6487.227 | 0.004 |
| Diglossa_brunneiventris | rump | slate | 1 | 4126.72  6 | 670.502 | 3456.22  4 | 0.187 | 3838.032 | 0.005 |
| Diglossa_sittoides | nape /  mantle | slate | 1 | 2704.34  9 | 797.101 | 1907.24  8 | 0.308 | 8310.673 | 0.002 |
| Diglossa_caerulescens | breast | slate | 3 | 1716.00  7 | 556.041 | 1159.96  6 | 0.325 | 36125.35  7 | 0.003 |
| Oreomanes_fraseri | nape | slate | 2 | 3978.62  9 | 824.841 | 3153.78  8 | 0.220 | 6779.868 | 0.002 |
| Conirostrum_cinereum | nape | slate | 2 | 2327.82  3 | 652.770 | 1675.05  3 | 0.280 | 27911.28  3 | 0.003 |
| Sporophila_schistacea | mantle | slate | 1 | 5113.27  3 | 953.833 | 4159.44  0 | 0.184 | 8103.644 | 0.004 |
| Tangara_vitriolina | mantle | slate | 10 | 1969.88  9 | 528.343 | 1441.54  6 | 0.268 | 67114.53  9 | 0.003 |

**Table S4.** Results of Phylogenetic generalized least square (PGLS) used to test the influence of predictor variables (nanostructure complexity, average hard block thickness, average soft block thickness, local crystallinity and I max) separately for PC1 of a) slate, b) blue and c) blue-slate-grey colours. Intercept estimates, standard error and p-values are reported.

|  | a) SLATE |  | b) BLUE | | c) BLUE-SLATE-GREY | |
| --- | --- | --- | --- | --- | --- | --- |
|  | Estimate (+/-  SE) | P  [anova] | Estimate (+/-  SE) | P  [anova] | Estimate (+/-  SE) | P [anova] |
| Intercept | 1.1975e-01 (+/-  7.4872e-02) |  | 5.3227e-01 (+/-  1.8853e+00) |  | 1.5104e-01 (+/- 3.6963e-  02) |  |
| nanostructure | 2.4639e-02 (+/-  1.8116e-02) | 0.25916 | 4.0498e-02 (+/- 1.6217e-  02) | 0.02315  * | 2.9624e-02 (+/- 7.9681e-  03) | 0.000895 |
| average_hard_block_thicnkess | 7.2865e-04 (+/-  2.6126e-04) | 0.42831 | 4.7362e-03 (+/- 3.3676e-  03) | 0.01042  * | 1.5021e-03 (+/- 3.2573e-  04) | 0.672532 |
| average_soft_block_thickness | -1.0313e-04  (+/- 4.1582e-  05) | 0.94045 | -7.5464e-04 (+/- 1.0810e-  03) | 0.08288 | -1.939e-04 (+/- 4.938e-  05) | 0.294091 |
| Local crystallinity | 1.3408e+00  (+/- 4.8115e-  01) | 0.013 | 8.088e+00 (+/-  7.658e+00) | 0.215 | 2.966e+00 (+/- 4.849e-  01) | 4.45E-08 |
| I max | -1.974e-06 (+/- 2.711e-06) | 0.481 | -1.1499e-06 (+/- 5.225e-  07) | 0.079 | 1.938e-06  (+/- 5.059e-  07) | 0.0005 |
| *Multiple R2* | 0.503 |  | 0.883 |  | 0.714 |  |
| *Adjusted R2* | 0.277 |  | 0.766 |  | 0.669 |  |
| *Lambda* | 0 |  | 1 |  | 0 |  |

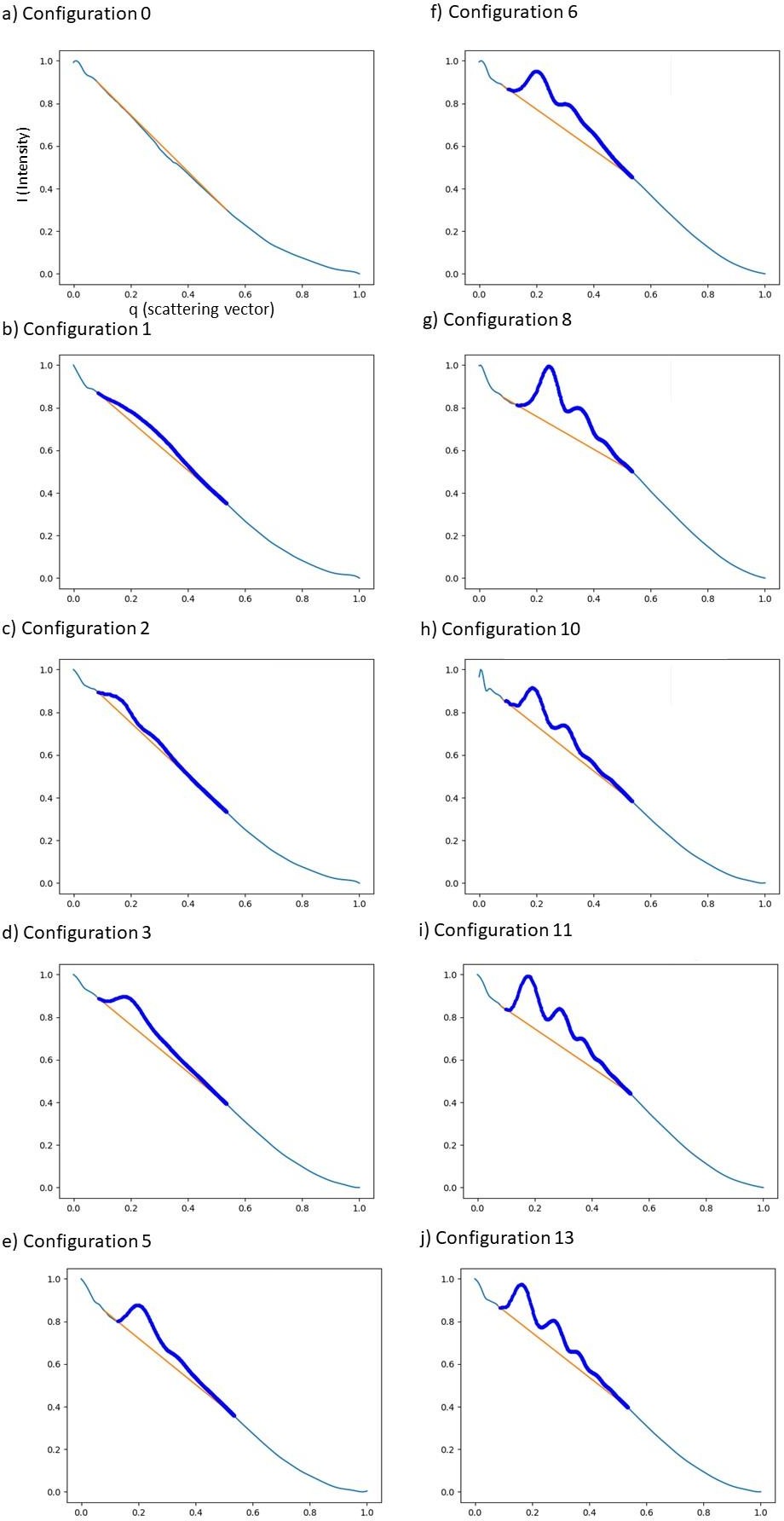

**Figure S1.** Examples of scattering profiles cover all possible peaks and shoulder combinations across all examined feather samples. This figure is the extension of the Figure 4.1. from the main text. The figure is a visual representation of Table 1, and the classification of combinations of features is explained in the table. In short, configuration 0 (a) indicates a lack of any structural components on the plot. Configurations 1 and 2 (b, c) have shoulder detected as the first scattering element and they are typical for the rudimentary form of the spongy nanostructure in the medullary feather cells. Configurations 3 and 5 (d, e) have peak detected as the first scattering element, and the following scattering element is either non-existing or a shoulder. Configurations 6, 8, 9, 10, 11 and 13 (f - j) have peaks detected as the first and second scattering element with any other number of elements (peaks or shoulders) detected afterwards. On each plot, an orange line represents an area that is explored for the detection of the peak and shoulders, thick blue line represents peak and shoulders detected, while a thin blue line represents all the data for each plot.
